# Human T Cells from Resistant Hypertension Patients Promote Hypertension via TNF in Humanized Mice

**DOI:** 10.64898/2026.01.23.701431

**Authors:** Masudur Rahman, Hendrik Bartolomaeus, Lydia Hering, Mina Yakoub, Guang Yang, Johannes Holle, Lukas Streese, Denada Arifaj, Milena Hahn, Marta Kantauskaite, Tim F. Grüner, Jaroslawna Meister, Nicola Wilck, Markus Kleinewietfeld, Lars C. Rump, Dominik N. Müller, Sebastian Temme, Johannes Stegbauer

## Abstract

**Objective:** Preclinical studies suggest a pivotal role of adaptive immunity, particularly T cells, in hypertension. However, due to the multifactorial pathogenesis, there is still no definitive evidence for a causal role of T cells in the development of hypertension in humans. We sought to determine whether T cells from patients with treatment-resistant hypertension (TRH) directly modulate blood pressure and vascular function *in vivo*.

**Methods:** Peripheral blood mononuclear cells (PBMCs) from TRH patients and healthy controls (HC) were adoptively transferred into immunodeficient NSG-(KbDb)^null mice. Hypertension was induced by angiotensin II infusion for 14 days and monitored continuously by radiotelemetry.

**Result:** Following T cell engraftment, blood pressure was assessed at baseline and during AngII infusion in both groups of recipient mice. At baseline, systolic blood pressure did not differ between both groups. However, mice receiving TRH-PBMCs developed a significantly higher systolic blood pressure following AngII compared with HC-PBMC recipients. Endothelial dysfunction in isolated perfused kidneys was more pronounced in AngII-challenged TRH-PBMC recipients compared to HC-PBMC recipients. TRH-PBMC recipients displayed elevated effector memory CD4⁺ T cells and Th17 frequencies in spleen and kidney, along with markedly increased renal expression of human T cell-derived TNF. Overnight incubation of mouse aortic rings with human TNF induced endothelial dysfunction, indicating a causal role of T cell-derived TNF. As a proof of concept, TNF inhibition attenuated AngII-induced hypertension in TRH-PBMC–engrafted mice.

**Conclusion:** T cells from patients with treatment-resistant hypertension promote an exaggerated hypertensive response and endothelial dysfunction in PBMC-engrafted humanized mice, promoted by TNF-mediated mechanisms. These findings provide evidence that T cell–derived TNF may contribute to the pathogenesis of human hypertension.

## Introduction

Despite the availability of multiple pharmacological therapies, hypertension remains a leading cause of cardiovascular morbidity and mortality globally, contributing to complications such as stroke, heart failure, and chronic kidney disease.^1,2^ Increasing evidence, particularly from animal studies, indicates that immune activation and chronic inflammation play a central role in the pathogenesis of hypertension, yet these mechanisms are not targeted by current treatments. Seminal work by Guzik et al. showed that mice deficient in adaptive immune cells are protected from hypertension, whereas adoptive transfer of T cells restores the hypertensive response to angiotensin (Ang)II.^3^ Among adaptive immune cells, activated pro-inflammatory T cell subsets - including effector memory T cells (TEM), T helper 1 (Th1) cells, and Th17 cells - play a particularly important role in modulating hypertension as well as vascular and renal injury.^4–7^ These T cell subsets secrete distinct cytokines, including tumor necrosis factor (TNF), interferon (IFN)-γ, and interleukin (IL)-17A, all of which are known to modulate vascular tone, function, and hypertensive renal remodeling.^8–10^

Elevated TNF levels have been found in hypertensive patients, including in CD8+ effector memory T cells re-expressing CD45RA (TEMRA) cells.^11^ Moreover, genetical deletion or pharmacological inhibition with etanercept reduces blood pressure and improves kidney function in hypertensive mice.^12,13^ TNF mediates these effects in part by decreasing nitric oxide (NO) bioavailability, suppressing eNOS expression, and vascular remodeling.^14–16^ IFN-γ, secreted by Th1 and CD8+ T cells, promotes fibrosis and glomerular filtration rate decline.^17^ Consistently, IFN-γ deficiency attenuates AngII induced blood pressure increase.^9^ IL-17A, produced predominantly by Th17 cells, promotes cardiovascular remodeling.^18,19^ IL-17A-deficient mice exhibit reduced blood pressures and are protected from hypertensive organ damage^10^, and IL-17A administration in mice induces hypertension and vascular remodeling.^20^ Similar to TNF, IFN-γ and IL-17A stimulate salt retention in the kidney and thereby elevate blood pressure.^21,22^

Among hypertensive individuals, those with treatment-resistant hypertension (TRH) are of particular concern, as TRH is associated with a dramatically increased risk of cardiovascular events and mortality.^23,24^ According to the guidelines of the European Society of Hypertension, TRH is defined as uncontrolled blood pressure despite the use of three adequately dosed antihypertensive agents.^25^ Notably, TRH is characterized not only by vascular dysfunction, increased sympathetic nerve activity (SNA), and renin-angiotensin-aldosterone system (RAAS) activation, but also by marked immune activation.^26–28^ Patients with TRH exhibit elevated plasma levels of C-reactive protein (CRP), TNF, and IFN-γ, along with a pro-inflammatory T cell signature characterized by higher proportions of effector memory CD4+ T cells.^11^

In the present study, we examined whether T cells from TRH patients exacerbate hypertension and explored the contribution of sustained pro-inflammatory cytokine secretion to hypertension and organ damage. To this end, we employed a humanized mouse model of hypertension by adoptively transplanting immunodeficient NSG-(KbDb)^null mice (NOD.Cg-Prkdc^scid^ H2-K1^tm1Bpe^ H2-D1^tm1Bpe^ Il2rg^tm1Wjl^/SzJ)^29^ with PBMCs from TRH patients (TRH-PBMC) or healthy controls (HC-PBMC). We experimentally induced hypertension by AngII infusion and subsequently analyzed inflammation and immune cell profiles in recipient mice from both groups. This unique approach enabled us to investigate the mechanisms by which T cells from TRH patients contribute to the development of hypertension and renal organ damage.

## Methods

The authors declare that all supporting data and analytical methods are available within the article and its online-only Data Supplement. The data, analytical methods, and study materials that support the findings of this study are available from the corresponding author upon reasonable request.

### Animal Ethics

Immunodeficient NSG-(KbDb)^null mice (JAX stock #023848) mice were used for the experiments. Animal experiments were carried out in accordance with the requirements of the Directive 2010/63/EU of the European Parliament on the protection of animals used for scientific purposes and approved by the federal state authority Landesamt für Natur-, Umwelt-, und Verbraucherschutz Nordrhein Westfalen (G338/16). Breeding and maintenance of the experimental mice were organized and conducted at the local animal care facility at Heinrich-Heine-Universität Düsseldorf. Mice were housed under standard conditions with free access to standard food and water ad libitum. Light to dark cycle was 12:12 hours. Mice were inspected daily for living environment and animation.

### Mouse strain and study design

A specialized immunodeficient mouse strain, NSG-(KbDb)^null, was chosen for the experiments because these mice lack mature T cells and B cells, do not have functional natural killer cells, and lack murine class I major histocompatibility complex (MHC) expression. As a result, the incidence of graft-versus-host disease is very low.^29^ Moreover, irradiation or implantation of human fetal lymphatic tissue is not necessary for successful engraftment of human immune cells. For experiments, male NSG-(KbDb)^null mice aged 6 to 8 weeks were used. Mice were supplied with neomycin (Sigma-Aldrich Chemie GmbH) in the drinking water throughout the experimental period. Peripheral blood mononuclear cells (PBMCs) were isolated from healthy controls and therapy-resistant hypertensive patients, and 2×10^6^ to 5×10^6^ PBMCs were transferred to each NSG-(KbDb)^null mouse. Three weeks after transfer, peripheral blood was collected from the mice to confirm successful engraftment by flow cytometry. After engraftment, telemetry catheters were implanted to measure blood pressure. Subsequently, AngII (500 ng/kg/min) osmotic minipumps (Alzet, California, USA) were implanted subcutaneously to induce hypertension. After 14 days of blood pressure measurement, mice were sacrificed for further assessment. Mice were anesthetized, and blood was withdrawn via the left renal artery, followed by euthanasia by cervical dislocation. Ice-cold PBS containing 100 U/mL heparin (B. Braun, Germany) was then injected into the left ventricle of the heart to flush remaining blood from the circulatory system, and organs were collected for further assessment.

### Study population

PBMCs from 8 male individuals with TRH (age>18) were collected. A secondary cause of hypertension was excluded and TRH was defined according to the guidelines of the European Society of Hypertension. Healthy controls are characterized as subjects without any cardiovascular diseases and a systolic blood pressure below 130 mmHg and absence of any antihypertensive drugs. All participants have signed the informed consent in accordance with the Declaration of Helsinki and approval from the Institutional ethics board of the University of Düsseldorf, Germany (Studiennummer 3848). Serum and human peripheral blood mononuclear cells (PBMC) were collected from patients with TRH and healthy controls and purified by Ficoll-Hypaque density centrifugation followed by cryopreservation in PBS containing 0.5% FCS (fetal calf serum) and 8% DMSO (Dimethyl sulfoxide). Later, frozen samples were thawed and washed for further analysis by flow cytometry or for injection into mice.

### Statistical Analysis

GraphPad Prism 10 was used for statistical analysis and graphical representation of the data. To compare two groups, a one- or two-tailed unpaired t-test (with or without Welch’s correction) or a Mann-Whitney test was used, if the data points did not fit to a Gaussian distribution. Two-tailed tests were conducted to investigate the hypothesis whether T cells from TRH affects AngII-dependent hypertension, the immune response and renal hypertensive organ injury. One-tailed tests were conducted to test the hypothesis whether T cells from TRH exaggerate cytokine expression and whether etanercept attenuates blood pressure response and the immune response. A two-way ANOVA test followed by Bonferroni multiple comparison was used to compare two independent variables between two groups. To obtain more statistically meaningful comparison, raw data were normalized in some experiments. Data were calculated as mean ± SEM. A *P* value <0.05 (*) was considered statistically significant.

## Results

### Human T cell engraftment in recipient mice enhances the hypertensive response

To investigate the role of human T cells in hypertension, we established a humanized mouse model in which human T cells can be engrafted without severe graft-versus-host disease (GvHD). For this purpose, we used immunodeficient NSG-(KbDb)^null mice as recipients. To engraft human T cells, 2×10^6^ to 5×10^6^ PBMC from healthy controls (HC) were injected intravenously (Fig-1A). Flow cytometry of peripheral blood after 3-4 weeks showed a sufficient abundance of human CD4+ (40.5±20.1 cells/µl) and CD8+ (8.90±3.99 cells/µl) T cells (Fig-1B and Suppl. Table 1), but no murine T cells (Fig-1B). No significant number of human monocytes, dendritic cells, B cells or natural killer (NK) cells were detected (Suppl. Fig-1A) indicating only successful engraftment of human T cells. Of note, after 2 weeks of AngII (500 ng/kg per min) infusion, flow cytometry demonstrated engraftment of human CD4+ and CD8+ T cells in the spleens and kidneys (Fig-1C). In contrast, no murine T cells were found in both spleens and kidneys of these mice (Fig-1C).

**Figure 1:**
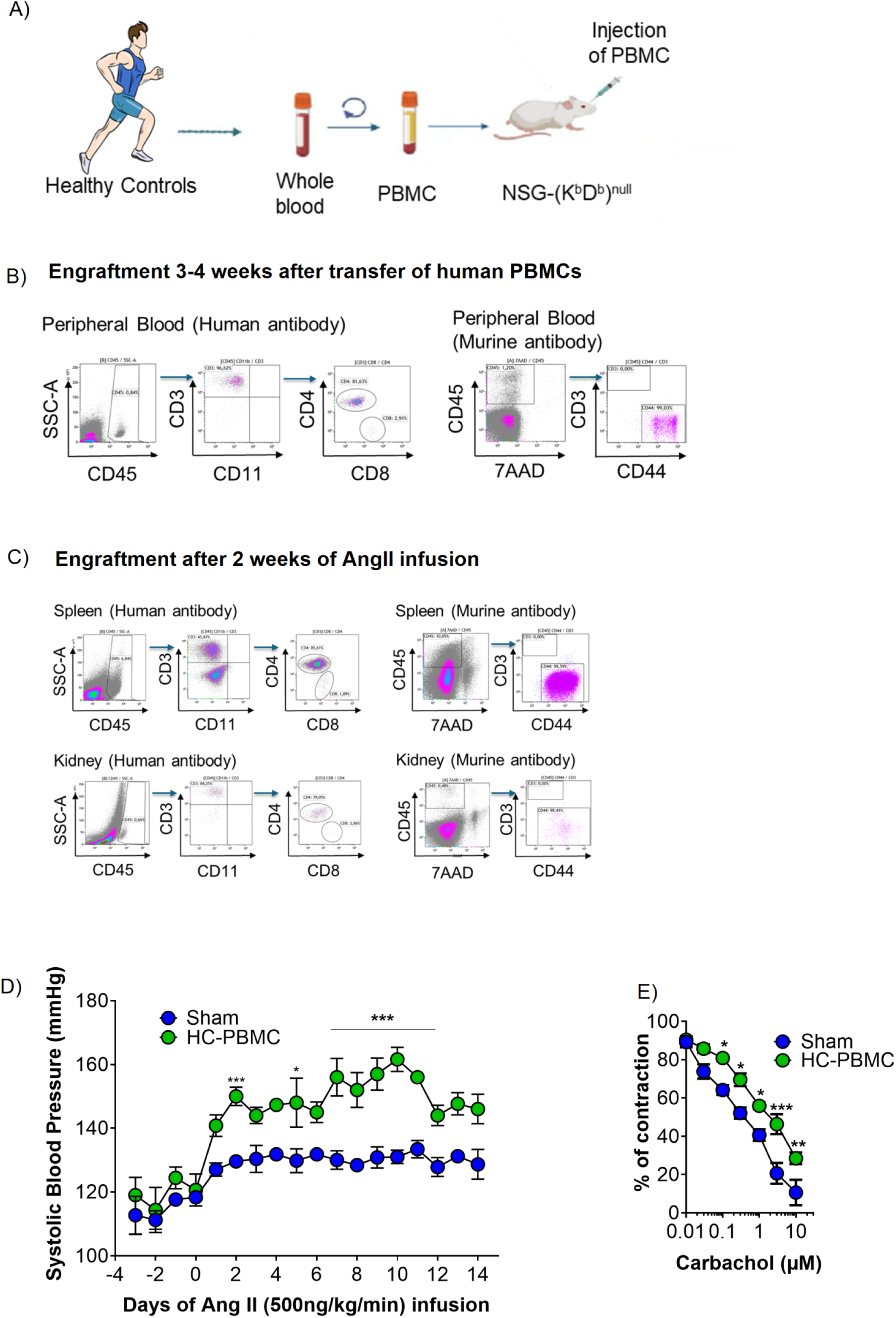
Successful human T cell engraftment in NSG-(KbDb)^null mice. (A) Pictogram of the experimental procedure. (B) Representative-gating pictures showed human CD3, CD4 and CD8 positive cells in peripheral blood of the mice but no murine CD3 positive cells after 3-4 weeks of human PBMC injection. (C) Representative gating showed presence of human CD3, CD4 and CD8 positive cells in the spleens and kidneys of mice infused with Ang II for 14 days but no murine CD3 positive cells. (D) Increased systolic BP response during chronic Ang II infusion in HC-PBMC mice (HC-PBMC versus Sham: baseline, 119 ± 2 versus 115 ± 2 mmHg, *P*>0.05, n=5; week 1, 147 ± 2 versus 130 ± 1 mmHg, *P*>0.05, n=5; week 2, 152 ± 2 versus 130 ± 1 mmHg, \**P*<0.05, n=5). (E) Endothelial-dependent renal vasorelaxation by carbachol was impaired in isolated perfused kidneys of HC-PBMC mice compared to Sham. Sham n=6, HC-PBMC n=6. \*\*\**P*<0.001, \*\**P*<0.01 \**P*<0.05, by two-way ANOVA followed by Bonferroni multiple comparison test was used to compare the difference between the groups from (D) to (E).

To examine whether human T cells were activated by AngII, relative expression level of various T cell-derived cytokines were measured in the kidneys by qPCR. Since only T cells engrafted and human-specific primers were used, measured expression levels reflect their production in T cells. Kidneys of successfully engrafted mice challenged with AngII show significantly increased mRNA expression levels of TNF, transforming growth factor (TGF)-ß, IL-2 and relatively higher expression of IFN-γ compared to the engrafted mice which were not treated with AngII (Suppl. Fig-1B). These results indicate that engrafted human T cells are activated by AngII *in vivo*, leading to increased production of pro-inflammatory cytokines.

To determine whether the human T cell transfer influences AngII dependent hypertension, systolic blood pressure was measured by tail cuff under baseline conditions and during chronic AngII infusion. At baseline, no difference in blood pressure was detected between mice engrafted with human T cells and mice without human T cells (119±2 versus 115±2 mmHg, *P*>0.05, n=5). After AngII infusion, blood pressure increased in both groups but was significantly higher in HC-PBMC (week 1: 147±2 vs. 130±1 mmHg, *P*>0.05; n=5; week 2: 152±2 vs. 130±1 mmHg, \**P*<0.05, n=5) (Fig-1D). In addition, endothelial dependent vasorelaxation of isolated perfused kidneys was significantly impaired in kidneys from AngII-infused HC-PBMC mice compared to control, indicating a driving effect of human T cells on renal vascular dysfunction (Fig-1E).

Taken together, human T cell engraft in NSG-(KbDb)^null mice without causing GvHD. Furthermore, the data demonstrates that our humanized model responds to AngII with an increase in BP and vascular damage.

### Pro-inflammatory T cell phenotype in patients with treatment resistant hypertension

Next, we recruited a cohort of normotensive healthy controls (HC) and patients with resistant hypertension to analyze the effects of hypertension-associated human T cell phenotypes in NSG-(KbDb)^null mice. We hypothesized that T cells from patients with TRH aggravate AngII-induced hypertension. Donor patients with TRH were slightly older and had significantly higher systolic and diastolic blood pressures. In this exploratory study, only male donors were used in order to reduce sex-related factors. Furthermore, patients with TRH took a median of 5 anti-hypertensive drugs (Table 1).

**Table 1:**
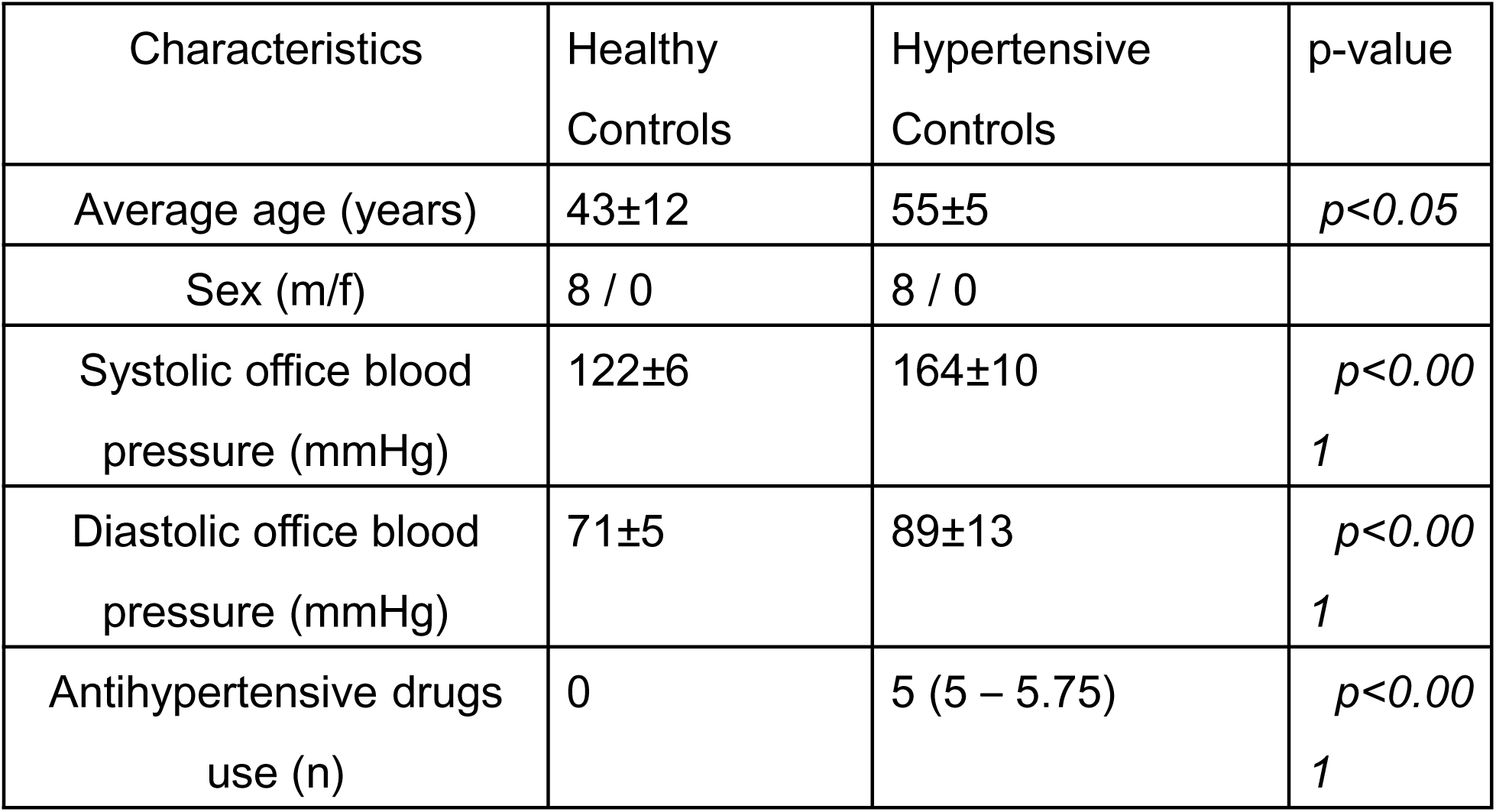
Baseline characteristics of PBMC donors.

To investigate T cell signatures which potentially affect the T cells response in engrafted mice, we analyzed PBMC by flow cytometry. Patients with TRH showed a significantly higher proportion of CD4+ TEM (CD45RA-CCR7-), CD4+ TEMRA (CD45RA+CCR7-) (Fig-2A), Th2 (CRTH2+CXCR3-) (Suppl. Fig-3A) and significantly lower proportion of CD4+ naïve T cells (TN, CD45RA+CCR7+) compared to healthy controls (Fig-2A). Proportions of CD4+ Th17 (CCR6+CXCR3-) (Fig-2A), central memory T cells (TCM, CD45RA-CCR7+), Th1 (CCR6-CXCR3+), regulatory T cells (Treg) (CD127-CD25+) were unaffected (Suppl. Fig-3A). For CD8+ T cells, patients with TRH showed increased CD8+ TCM compared to healthy controls (Fig-2B), while no difference was observed in the proportion of CD8+ TEM, TEMRA and TN (Suppl. Fig-3B).

**Figure 2:**
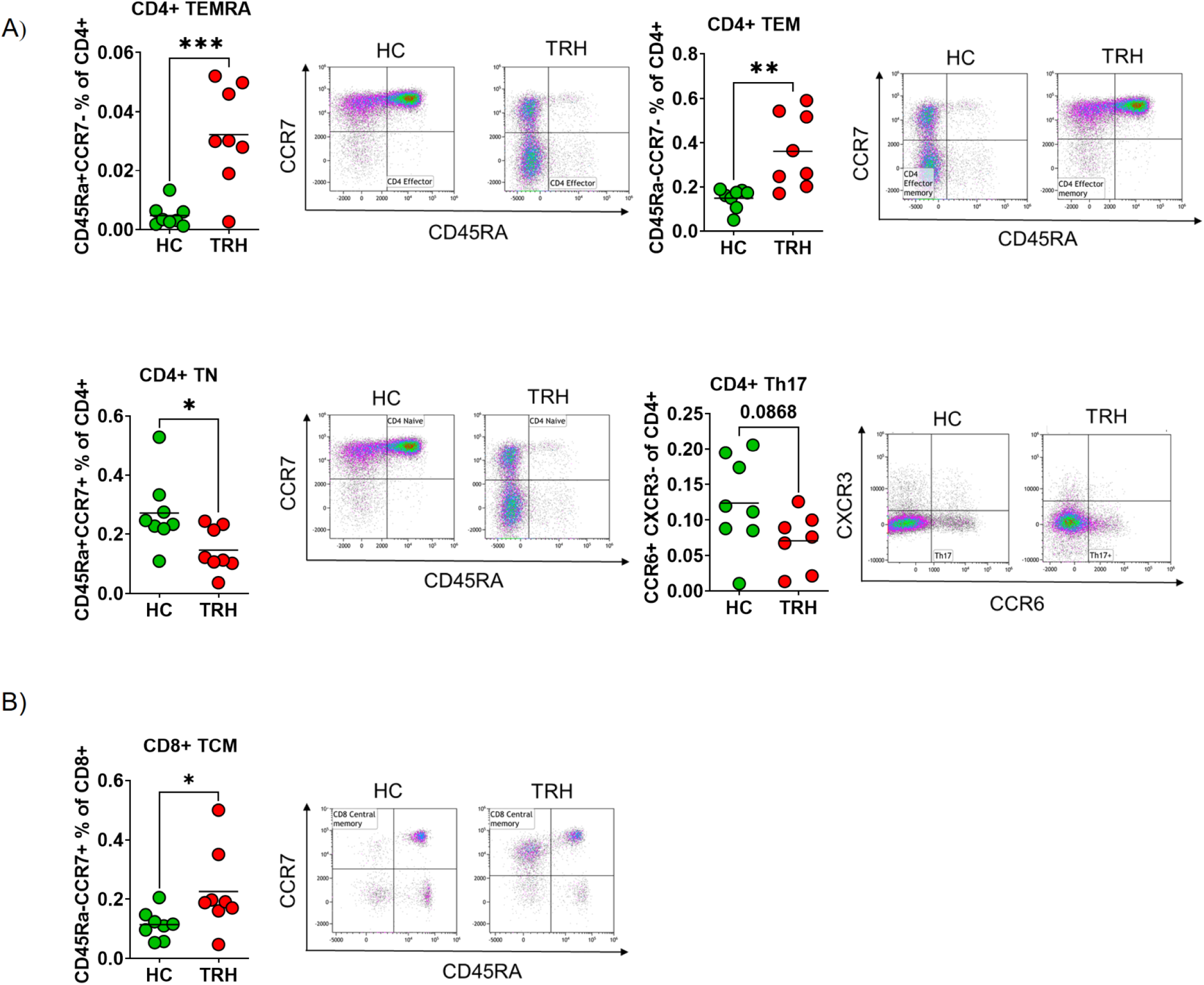
Pro-inflammatory T cell signatures of healthy and treatment resistant hypertensive donors. (A) PBMCs were analysed for CD4+ TEM (CD45Ra-CCR7-), TEMRA (CD45Ra+CCR7), TN (CD45Ra+CCR7+), Th17 (CCR6+CXCR3-), cells. Left, quantification in percentage of CD4+ cells. Right, representative plot of flow cytometry. HC n=8, TRH n= 5-8. (B) PBMCs were analysed for CD8+ TCM (CD45Ra-CCR7+), cells. Left, quantification in percentage of CD8+ cells. Right, representative plot of flow cytometry. HC n=8, TRH n= 5-8. \*\*\**P*<0.001, \*\**P*<0.01 \**P*<0.05, by 2-tailed unpaired t-test.

In summary, these data confirm that patients with TRH show an altered T cell phenotype with an increase in antigen experienced T cell subsets like CD4+ TEM and CD4+ TEMRA.

### Renal inflammation is amplified in mice engrafted with T cells from TRH patients

To investigate whether T cells from TRH patients cause an altered pro-inflammatory T cell phenotype in response to AngII in mice, NSG-(KbDb)^null mice were engrafted with T cells from either TRH patients or healthy controls (HC). Splenic and kidney-infiltrating immune cells were analyzed by flow cytometry after 14 days of AngII infusion. Kidney tissue from TRH-PBMC transplanted mice contained significantly increased proportions of CD4+ TEM and Th17 cells compared to HC-PBMC mice, whereas proportions of CD4+ TCM were significantly reduced (Fig-3A). Similar effects on T cell subsets were observed in the spleen (Suppl. Fig-5A & 5B). CD4+ TN, TEMRA, Th1, Th2, and Treg were unaffected in the kidneys (Suppl. Fig-4A & 4B) as well as in the spleens (Suppl. Fig-5A & 5B). Moreover, CD8+ TEM frequencies in the kidney were significantly higher, whereas the TCM cell population was significantly reduced in TRH-PBMC mice compared to HC-PBMC mice (Suppl. Fig-4A & 4C). In contrast, splenic CD8+ TEM and TCM were not affected (Suppl. Fig-5A & 5C). Additionally, no difference was found in the populations of CD8+ TN and TEMRA in the kidneys (Suppl. Fig-4A & 4C) and spleens (Suppl. Fig-5A & 5C) of TRH-PBMC and HC-PBMC mice. These data show only a partial overlap with the T cell phenotypes prior to transfer, with CD4+ TEM remaining increased in AngII-infused mice.

**Figure 3:**
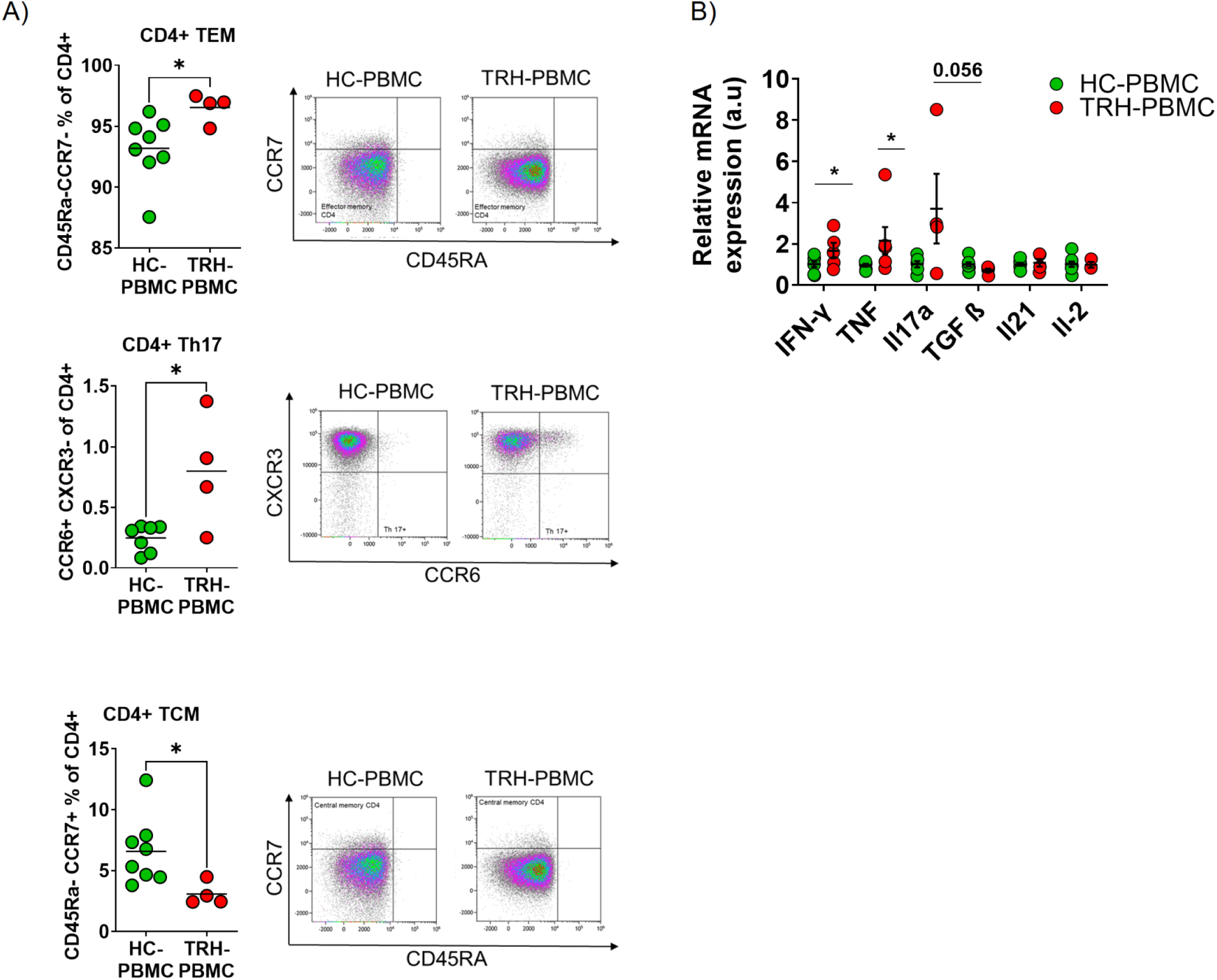
Increased pro-inflammatory human T cell subsets in kidneys of TRH-PBMC mice. (A) Flow cytometry analysis of kidney cells showed higher proportions of CD4+ TEM (CD45Ra-CCR7-), lower proportions of TCM (CD45Ra-CCR7+) and higher proportions of Th17 (CCR6+CXCR3-) cells in the kidneys of TRH-PBMC compared to HC-PBMC mice. HC-PBMC n=7-8, TRH-PBMC=4. (B) Increased human T cell derived pro-inflammatory cytokines such as TNF-α, IFN-γ, in the kidney tissue of TRH-PBMC compared to HC PBMC mice. l. HC-PBMC n=6-8, TRH-PBMC n=3-6. \**P*<0.05, by one tailed unpaired t-test.

Since pro-inflammatory engrafted human CD4+ TEM and Th17 were more abundant in kidneys and spleens of TRH-PBMC mice, we hypothesized that these T cell subsets are the main contributors to pro-inflammatory cytokine secretion, hypertension, and renal damage. To test this hypothesis, expression of pro-inflammatory T cell-derived cytokines in the kidney were assessed. Relative expression of human pro-inflammatory cytokines such as *TNF* (2.15±0.67 vs. 1.00±0.07 n=5-6, \**P*<0.05) and *IFN-γ* (1.67±0.38 vs. 1.00±0.15; n=5-8, \**P*<0.05 were higher in kidneys of TRH-PBMC mice compared to HC-PBMC mice (Fig-3B). No differences for *IL-17A, IL-2*, *TGF-ß*, and *IL-21* were observed between the two groups (Fig-3B). Of note, for murine cytokine isoforms we found either no detectable expression or non-significant differences between the two groups (Suppl. Fig-7). These results suggest that T cells from TRH-PBMC mice release more pro-inflammatory *TNF and IFN-γ* than T cells from HC-PBMC mice and thereby promote hypertension and organ damage.

### Mice engrafted with T cells from TRH patients exhibit increased hypertension and renal damage

To investigate whether the amplified pro-inflammatory cytokine response of engrafted human T cells from TRH patients affect the blood pressure response to AngII and kidney injury, NSG-(KbDb)^null mice were engrafted with either T cells from patients (TRH-PBMC) or T cells from healthy controls (HC-PBMC). Blood pressure was measured continuously by radiotelemetry under baseline condition and during 14 days of continuous AngII infusion (Fig-4A). Under baseline conditions, no difference in systolic blood pressure was found between TRH-PBMC and HC-PBMC mice (130±2 vs. 131±2 mmHg, *P*>0.05, n=6). During AngII infusion, TRH-PBMC mice showed significantly higher systolic blood pressures compared to HC-PBMC mice, with the strongest effect during the first week (week 1, 161±1 vs. 135±2 mmHg, \**P*<0.05, n=6); week 2, 156±2 vs. 140±2 mmHg, *P*>0.05, n=6) (Fig-4B and Suppl. Fig-6). In line, hypertensive kidney damage was significantly exacerbated in AngII-infused TRH-PBMC mice compared to HC-PBMC mice. Plasma cystatin C levels indicated a lower glomerular filtration rate (1347±167 vs. 945±77 ng/ml, \**P*<0.05, n=6-7) (Fig-4D) in TRH-PBMC mice. Renal *ex vivo* endothelial dependent vasodilation was significantly impaired in TRH-PBMC mice (Fig-4C). Moreover, renal vascular wall hypertrophy assessed by vessel wall to lumen ratio (4.11±0.06 vs. 3.59±0.13 A.U, \**P*<0.05, n=5-7) as well as perivascular immune cell infiltration was significantly aggravated in TRH-PBMC mice (Fig-4E-F).

**Figure 4:**
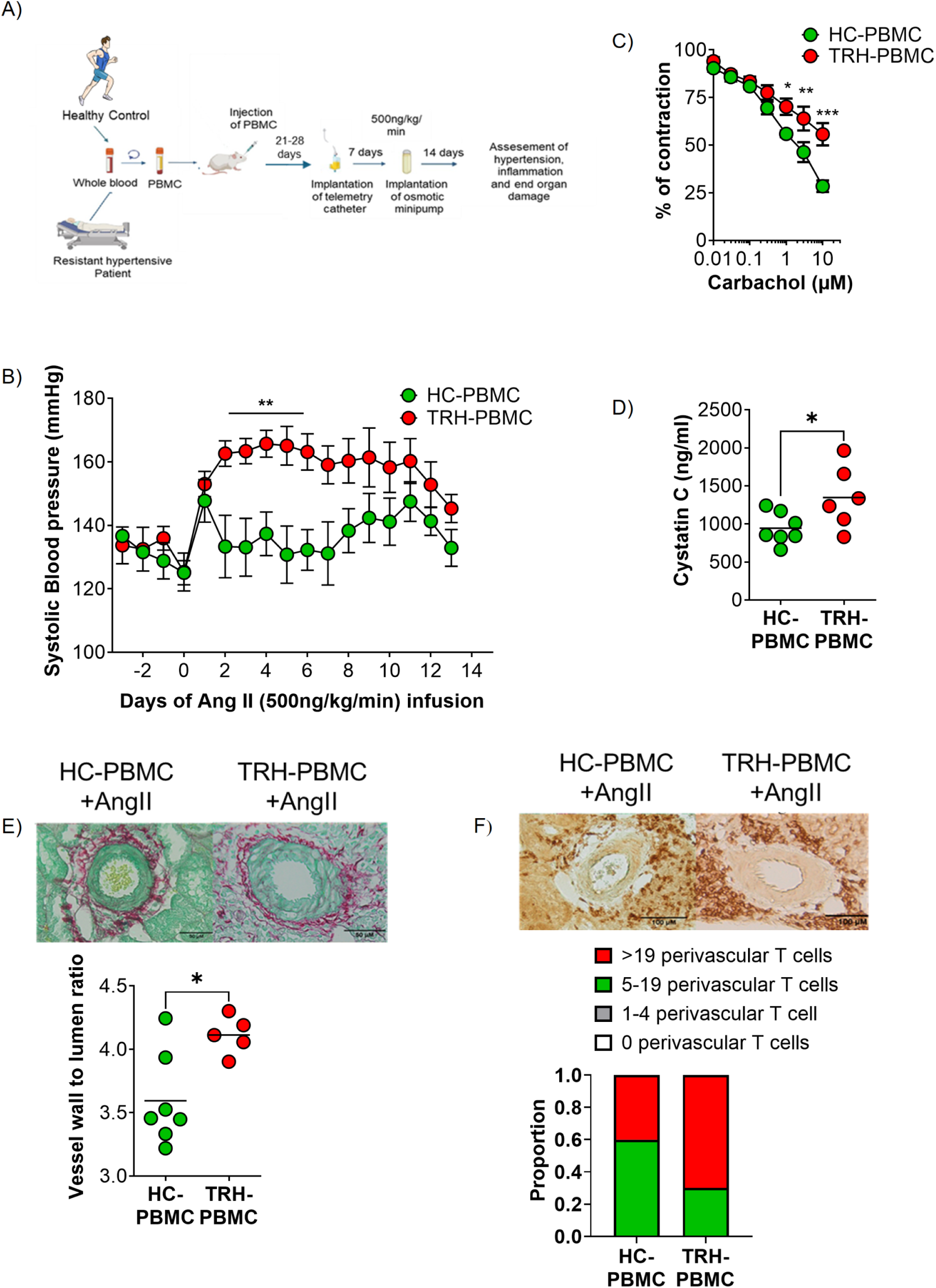
Exaggerated blood pressure response and renal damage during chronic Ang II infusion in TRH-PBMC mice. (A) Pictogram diagram of the complete experimental procedure. (B) Increased systolic blood pressure response in TRH-PBMC compared to HC-PBMC mice after AngII infusion (TRH-PBMC versus HC-PBMC: baseline, 130 ± 2 versus 131 ± 2 mmHg, *P*>0.05, HC-PBMC n=6, TRH-PBMC n =6; week 1, 161 ±1 versus 135 ± 2 mmHg, \*\**P*<0.01, HC-PBMC n=6, TRH-PBMC n =6; week 2, 156 ± 2 versus 140 ± 2 mmHg, *P*>0.05, HC-PBMC n=6, TRH-PBMC n =6). (C) Endothelial-dependent renal vasorelaxation by carbachol was impaired in isolated perfused kidneys of TRH-PBMC mice compared to HC-PBMC mice. HC-PBMC n=5, TRH-PBMC n=5. \*\*\**P*<0.001, \*\**P*<0.01, \**P*<0.05. (D) Plasma cystatin C was elevated in TRH-PBMC mice compared to HC-PBMC mice (1347± 167 versus 945 ± 77 ng/ml, \**P*<0.05, HC-PBMC n=7, TRH-PBMC n =6. (E) Renal vascular hypertrophy determined by wall to lumen ratio was higher in TRH-PBMC mice compared to HC-PBMC mice (4.11 ± 0.06 versus 3.59 ± 0.13 A.U, \**P*<0.05). HC-PBMC n=7, TRH-PBMC n =5. (F) Increased infiltration of CD3 positive cells in the perivascular regions of the kidneys of the TRH-PBMC mice compared to HC-PBMC mice. Two-tailed unpaired t-test or Two-way ANOVA followed by Bonferroni multiple comparison test was used to compare the difference between the groups from (B) to (E).

Taken together, human T cells from TRH donors elicit an increased hypertensive response compared to HC-derived T cells along with an exacerbated renal damage and vascular remodeling.

### TNF induces endothelial dysfunction and promotes AngII-dependent hypertension

To test whether human TNF and IFN-γ exert their proinflammatory effects in mouse kidney tissue, we first incubated kidney slices with either TNF or IFN-γ and measured the expression of damage-associated target genes (Suppl. Fig.-8A). IFN-γ (100 U/mL) did not increase the expression of its target gene (Suppl. Fig-8C) but interestingly, TNF (0.1 ng/mL) increased the expression of adhesion molecules *Vcam1* and *Icam1* (Suppl. Fig-8B). Based on these results, we tested the effect of human TNF on murine vascular function. Therefore, isolated murine thoracic aortic rings were incubated with TNF (4 nM) for 22 hours, after which endothelial function was tested by wire myography. In norepinephrine pre-constricted aortas, carbachol-induced endothelium-dependent vasorelaxation was significantly impaired after TNF incubation, indicating an important role of human TNF in vascular injury (Fig-5A). To test whether *in vivo* inhibition of human TNF improves vascular function and attenuates exacerbated AngII-induced hypertension, TRH-PBMC mice were injected with the clinically used TNF decoy receptor etanercept (0.8 mg/KG) and compared against mice that received a control injection with solvent. Under baseline conditions, no difference in systolic blood pressure (136±2 versus 141±1 mmHg, *P*>0.05, n=6) was observed. During continuous AngII infusion, blood pressure was significantly higher in solvent treated than in the etanercept-treated mice (week 1: 159±4 vs 136±2 mmHg, *P*>0.05, n=6; week 2: 161±1 vs 130±2 mmHg, \**P*<0.05, n=6) (Fig-5B). To check whether etanercept affects T cell subsets, flow cytometry analysis of spleen and kidney-infiltrating leukocytes was performed. Spleens of solvent treated mice showed significantly increased proportions of CD4+ TEM compared to etanercept-treated mice, whereas CD4+ Treg frequencies were significantly reduced (Suppl. Fig-9A). On the contrary, CD4+ TN, TEMRA, TH17, Th1, Th2, and TCM were unaffected in the spleen (Suppl.Fig-9A). Furthermore, CD4+ TEM, TN, TEMRA, TCM, Th1, Th2, Th17 and Treg (Suppl.Fig-10A) as well as CD8+ TEM, TN, TEMRA, TCM remained unchanged in the kidneys (Suppl.Fig-10B).

**Figure 5:**
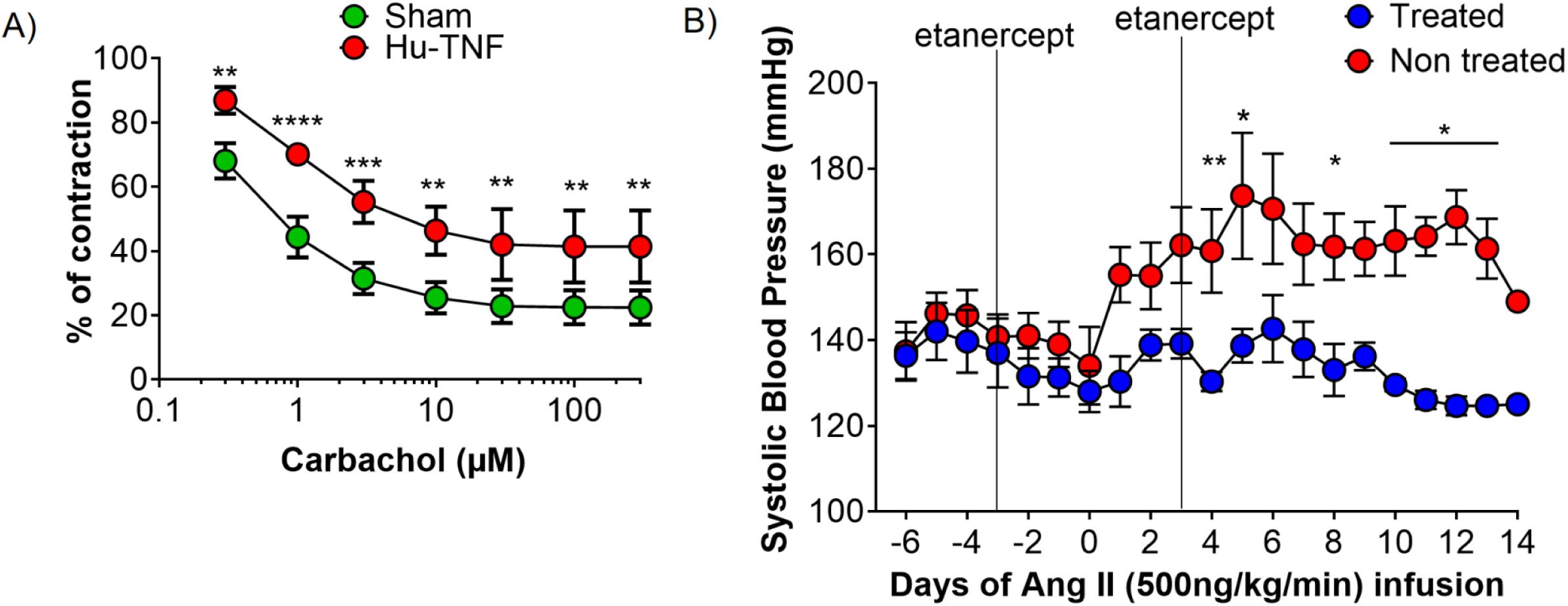
TNF induces endothelial dysfunction and promotes AngII-dependent hypertension. (A) Overnight incubation of mouse aortic rings with human TNF induced endothelial dysfunction. Endothelial-dependent vasorelaxation was expressed as percentage of contraction induced by norepinephrine. Sham n=7, Human TNF n=4. \**P*<0.05, \*\**P*<0.01, \*\*\**P*<0.001. (B) Inhibition of TNF attenuates systolic blood pressure response in TRH-PBMC mice (treated versus non-treated: baseline, 136 ± 2 versus 141 ± 1 mmHg, *P*>0.05, Treated n=7, non-treated n=6; week=1, 136 ± 2 versus 159 ± 4 mmHg, *P*>0.05, Treated n=6, non-treated n =6; week 2, 130 ± 2 versus 163 ± 1 mmHg, \**P*<0.05, Treated n=6, non-treated n =6). Two-way ANOVA followed by Bonferroni multiple comparison test was used to compare the difference between the groups.

In summary, we demonstrate that the enhanced T cell-mediated TNF production observed in TRH patients is associated with renal and vascular damage following transfer of TRH-derived T cells into mice, underscoring the relevance of T cell-derived TNF in human hypertension.

## Discussion

Preclinical studies have demonstrated the influence of particularly T cell-mediated immunity, on hypertension and renal damage. However, translational evidence demonstrating the relevance of human T cells for hypertension and associated organ damage is lacking. Here, we provide evidence for the mechanistic relevance of T cell-driven inflammation in TRH and hypertensive renal damage. In this study, we aimed to investigate the contribution of human T cells to hypertension and renal injury. First, we established a humanized mouse model that enabled us to determine the specific functions of human T cells under hypertensive conditions. Second, we demonstrated that T cells, particularly CD4+ TEM and Th17 cells from TRH human donors, enhance AngII-induced hypertension in humanized recipient mice, alongside aggravated kidney damage and inflammation. Third, we show that T cell-derived TNF is a key to these deleterious hypertensive effects.

Research on the role of human T cells in the development of hypertension is limited by ethical and practical constraints. As a result, much of the existing data comes from *in vitro* models that insufficiently reflect the complexity of the interaction of human immune responses with the multi-layered facets of blood pressure regulation. To overcome this limitation, we employed a specialized immunodeficient mouse strain, NSG-(KbDb)^null.^29^ This strain offers several advantages for our study. First, engraftment with 10×10⁶ human PBMC leads to robust human T cell reconstitution. Second, neither harmful irradiation nor implantation of human fetal lymphoid tissue is required. Third, the incidence of GvHD is low and typically occurs earliest 16 weeks post-engraftment.^30^ We thus successfully engrafted PBMC-derived T cells in these mice. Furthermore, after two weeks of AngII infusion, a significant infiltration of human donor CD4⁺ and CD8⁺ T cells in spleens and kidneys became apparent. Notably, we did not detect significant amounts of engrafted human non-T immune cell populations (CD45+ CD3-), nor relevant responses of murine T cells in the blood, spleen, or kidneys, indicating that a dominant role of the engrafted human T cells in our model. The altered T cell phenotype found in patients with TRH was only partially transferred into NSG-(KbDb)^null mice, likely related to model-related constraints such as the xenogeneic nature of the transplantation as well as the experimental induction of hypertension by AngII infusion. The specific engraftment of T cells in humanized mice using PBMCs was expected based on previous studies^31,32^, whereas successful engraftment of myeloid populations typically requires transplantation of CD34⁺ human hematopoietic stem cells derived from fetal liver tissue.^33^ In contrast to our approach, Itani et al. used such humanized mice to study the effects of experimental hypertension on human T cells.^34^ Because our study focused on T-cell phenotypes derived directly from patients with TRH, this approach was not feasible. Importantly, this distinction highlights an advantage of our proposed model system, which allows investigation of the functional impact of distinct human T-cell populations. This approach may also be extended to study other cardiovascular risk factors associated with altered T-cell phenotypes in chronic kidney disease.

In this study, we examined T cell phenotypes associated with TRH. Hypertension is generally linked to a pro-inflammatory shift in the immune system, particularly among T cells. In human hypertension, lower frequencies of regulatory T cells (Treg) and exhausted CD8+ T cells have been reported^35^ along with increased Th1 and Th17 responses and higher frequencies of CD8+ and CD4+ TEM and TEMRA cells.^11,36,37^ The T cell profile observed in our TRH patient cohort partially overlaps with these findings. Notably, we also observed increased frequencies of Th2 cells using specific surface markers CRTH2+CXCR3-. It should be noted that our cohort is limited by its small sample size, as it was primarily designed for PBMC transfer studies into NSG-(KbDb)^null mice.

To further characterize our humanized mouse model, we analyzed the cytokine production of the engrafted human T cells in response to Ang II-induced hypertension. Since only human T cells were successfully engrafted, tissue cytokine mRNA expression is primarily explained by the engrafted T cells. We observed increased expression of human cytokines in the kidneys of Ang II–infused hypertensive compared to non-hypertensive engrafted mice. We observed an increase in *TNF*, *TGF-ß*, *IL-2* and a trend for *IFN-γ*. These findings indicate that hypertension is associated with an increased cytokine response by human T cells in target organs such as the kidney. Of note, TNF plasma levels have been independently associated with blood pressure in hypertensive individuals.^38^ Previously, we have confirmed this in patients with TRH and demonstrated increased production of TNF and IFN-γ by TEMRA.^11^ TNF and IFN-γ plasma levels are associated with incident hypertension.^39^ While there are no reports on IL-2 in hypertensive patients, TGF*-ß* has been associated with hypertension risk^40^ as well as established hypertension.^41^ Interestingly, we observed no significant increase in IL-17A despite *in vitro* associations with AngII^42^ and a strong association with CVD.^43^ In general, we found that Ang II–infused mice engrafted with human T cells exhibited a significantly greater blood pressure increase and more pronounced impairment of renal endothelial vasodilation compared to non-engrafted mice. These results are consistent with the literature on both T cell deficient mouse models as well as immune system humanized mice.^3^ Taken together, our results demonstrate the successful establishment of a humanized mouse model in which human T cells not only engraft robustly, infiltrate target organs and respond to AngII, thus providing a relevant *in vivo* platform to study human T cell-mediated mechanisms in hypertension.

The main goal of this study was to investigate the specific contribution of human T cells from individuals with TRH. Mice receiving TRH-derived T cells developed significantly higher blood pressures and more pronounced signs of hypertensive kidney damage compared to controls. Notably, engrafted T cells predominantly infiltrated the perivascular space of the kidney, a distribution associated with local cytokine release and vascular injury ^44^ which may contribute to increased vascular stiffness, endothelial dysfunction, and vascular hypertrophy.^13,45,46^ Consistent with this, we observed significantly greater renal vascular hypertrophy and endothelial dysfunction in the kidney vasculature of TRH-PBMC mice. However, our study cannot determine whether this vascular remodeling is primarily caused by the excessive blood pressure response or by proinflammatory signals from T cells. It is important to note that endothelial dysfunction is a hallmark of hypertension.^47–49^

To better understand the mechanism underlying the aggravated hypertensive phenotype in mice engrafted with T cells from individuals with TRH, we analyzed T cell subset distribution and pro-inflammatory cytokine expression in the spleen and kidney. Our data revealed a significantly increased presence of human TEM both CD4⁺ and CD8⁺ in the kidneys and spleens of TRH-PBMC mice compared to HC-PBMC mice. TEM cells are a differentiated subset of memory T cells characterized by rapid effector function upon restimulation.^50^ They are known to circulate in blood and peripheral tissues and to produce pro-inflammatory cytokines such as TNF and IFN-γ in response to antigenic or inflammatory cues^50,51^ and play a substantial role in the development of cardiovascular disease.^6,52–54^ Furthermore, we observed an increase in Th17 cells in the kidneys of TRH-PBMC mice compared to HC-PBMC mice. Consistent with these cell types we observed an increased expression of TNF and IFN-γ as well as strong trends for IL-17A in the kidney of TRH-PBMC mice. Of note, TNF and IFN-γ showed an increase in response to AngII in mice with T cells from healthy donors. In line with the extensive literature on these cytokines, we hypothesized a causal role for these cytokines for the aggravated kidney injury in mice with T cells from TRH patients. We focussed with our *in vitro* approaches on TNF and IFN-γ. TNF impaired vascular function *in vitro* as well as led to an upregulation of adhesion molecules in the kidney. In line, *in vivo* TNF blockade using etanercept ameliorated the hypertensive phenotype in TRH-PBMC mice. The mechanistic link between TNF and endothelial dysfunction is well established in both animal and human studies.^16,55^ TNF promotes oxidative stress, reduces nitric oxide (NO) bioavailability, and impairs endothelium-dependent vasodilation.^16,56,57^ These findings are supported by previous findings demonstrating increased TNF levels in hypertensive patients^38,58^ and the protective effects of TNF inhibition^59^ or deletion^60^ in experimental hypertension. Our findings extend this knowledge and show that the mechanism described in mice are similarly important in T cells from hypertensive patients.

Taken together, these results demonstrate that human TNF, produced by pro-inflammatory T cells from TRH individuals, is biologically active in murine tissues and may contribute to blood pressure elevation and endothelial dysfunction. Thus, TNF may represent a promising therapeutic target in treatment-resistant hypertension.

## Limitation

In the current study, we developed a humanized mouse model to investigate the role of human T cells in hypertension and hypertensive kidney damage. Although the model is based on the transfer of human PBMCs, we observed selective proliferation of human T cells, whereas antigen-presenting cells did not persist. In addition, T cells are likely activated through contact with xenogeneic murine tissue, such that physiological interactions between human T cells and antigen-presenting cells are not fully recapitulated. Moreover, certain effects may be underestimated, as human-derived cytokines do not activate murine receptors, as described for IFN-γ. Despite these limitations, this model provides a robust platform to study the effects of activated and proinflammatory human T cells under pathological conditions such as hypertension. Accordingly, it represents a suitable PBMC-based humanized mouse model for cardiovascular research.

### What Are the Clinical Implications?

- PBMC based humanized mouse model to investigate effects and impact of T cells from treatment resistant hypertension.
- Pro-inflammatory T cell signature from therapy resistant hypertension aggravates blood pressure response and renal vascular dysfunction.
- TNF released by human T cells is associated with exaggerated blood pressure and inhibition of TNF attenuates the blood pressure response to Ang II in TRH-PBMC mice. Modulating TNF release could be a promising approach to improving blood pressure response in individuals with treatment-resistant hypertension.
- This platform is suitable to investigate the contribution of altered T cell phenotypes found with other cardiovascular risk factors

## Acknowledgments

The authors thank Blanka Duvnjak, Christina Schwandt and Nicol Kuhr for their excellent assistance. J.S., L.H. and M.R. led and conceived the project, and designed the experiments. H.B., J.H., S.T., M.H. and M.K. isolated immune cells from humans and mice and performed flow cytometry analysis. R.M. L.H., M.Y., G.Y., D.A., T.F.G. L.S. and J.M. performed mice experiments analyzing blood pressure, vascular as well as kidney function and damage. D.N.N., M.K., N.W. and L.C.R gave major conceptional input. R.M. and J.S. wrote the paper with key editing by S.T., N.W., H.B., and J.M. and further input from all authors. The authors also thank BioRender (biorender.com) for providing items for drawing scheme graphs.

